# The Sustained Alteration Of Brain Waves In Cynomolgus Macaques Following Aerosol Infection With Venezuelan Equine Encephalitis Virus Subtype IAB

**DOI:** 10.64898/2026.08.28.747802

**Authors:** Sara I. Ruiz, Michael V. Accardi, Franco D. Rossi, Stephanie V. Trefry, Thomas R. Sprague, Joshua D. Shamblin, April M. Babka, Jun Liu, Xiankun Zeng, John C. Trefry, Simon Authier, Margaret L. Pitt, Farooq Nasar

## Abstract

Venezuelan equine encephalitis virus subtype IAB (VEEV-IAB) is a mosquito-borne virus that can cause fatal encephalitis in humans and equids. During the 20th century, sporadic but widespread outbreaks occurred throughout the Americas. In addition, VEEV-IAB was investigated as a potential biological warfare agent during the Cold War. Currently, no countermeasures are available to treat or prevent human infection. A critical impediment to understanding VEEV-IAB pathogenesis and developing countermeasures is the lack of a detailed disease course in a susceptible animal model. This study evaluated VEEV-IAB disease progression in cynomolgus macaques using advanced telemetry technology to continuously monitor physiological parameters, including temperature, respiration, activity, heart rate, blood pressure, electrocardiography (ECG), and electroencephalography (EEG), following an aerosol challenge of 6.0 log_10_ PFU. Following infection, all parameters were altered relative to baseline; temperature (+3.1 to +4.0°C), respiration rate (+45 to +91%), activity [daytime (- 29 to -55%) and nighttime (+14 to +34%)], heart rate (-27 to +191%), systolic (+11 to +39%) and diastolic blood pressure (+7 to +39%). Cardiac abnormalities included increases in QTc (Bazett), PR interval, and QRS duration. All EEG frequency bands were rapidly altered (−250% to +4,800%) and did not return to baseline during the 28-day post-infection period. Despite these profound physiological changes, brain tissues collected at 28 dpi showed minimal evidence of viral persistence or pathology. These data demonstrate that VEEV-IAB aerosol infection rapidly and markedly alters physiological parameters regulated by the autonomic nervous system, as well as provides new insights into VEEV-IAB pathogenesis and countermeasure development.

**IMPORTANCE:** VEEV-IAB is a high-consequence arbovirus that can cause fatal neurological disease, however, currently there are no approved vaccines or therapeutics. In this proof-of-concept study, we utilized state-of-the-art telemetry technology to characterize the disease course following VEEV-IAB infection in a susceptible macaque model by measuring multiple physiological parameters relevant to human disease. VEEV-IAB infection rapidly induces substantial alterations in autonomic nervous system functions including temperature, respiration, activity, heart rate, blood pressure, ECG, and EEG. Most notable findings were the extreme and sustained alterations of brain activity despite pathological changes in the brain. These findings establish a framework for disease assessment through quantification of critical physiological biomarkers and provide a platform for evaluating the efficacy of vaccine and therapeutic candidates against VEEV-IAB.

## INTRODUCTION

The genus *Alphavirus,* in the family *Togaviridae,* is comprised of spherical, enveloped viruses with single-stranded, positive-sense RNA genomes, ∼11-12 kb in length (1, 2). The genus consists of 32 recognized species, classified into eleven complexes based on antigenic and genetic similarities. The two aquatic alphavirus complexes [Salmon pancreatic disease virus (SPDV) and Southern elephant seal virus (SESV)] are not known to utilize arthropods in their transmission cycles, whereas all of the remaining complexes [Barmah Forest, Ndumu, Middelburg, Semliki Forest, Venezuelan (VEE), eastern (EEE), western equine encephalitis (WEE), Trocara, and Eilat] consist of arboviruses that almost exclusively use mosquitoes as vectors (1–6). Mosquito-borne alphaviruses infect diverse vertebrate hosts, including equids, birds, amphibians, reptiles, rodents, pigs, nonhuman primates, and humans (1). Human infections with Old World alphaviruses, such as chikungunya, o’nyong-nyong, Sindbis, and Ross River, are rarely fatal and are characterized by rash and debilitating arthralgia that can persist for months or years. In contrast, New World alphaviruses, such as eastern (EEEV), western (WEEV), and Venezuelan equine encephalitis virus (VEEV), can cause fatal encephalitis.

Members of the Venezuelan equine encephalitis (VEE) complex are important human and veterinary pathogens in the Americas as they can cause fatal disease in both humans and equids. The complex is comprised of thirteen viruses, including six subtypes and seven varieties. The first member of the complex, VEEV subtype IAB, was isolated in 1938 in Yaracuy State, Venezuela (7, 8). Following its discovery, multiple members of the complex [(IC, ID, IE, IF (Mosso das Pedras), II (Everglades), IIIA (Mucambo), IIIB (Tonate), IIIC, IIID, IV (Pixuna), V (Cabassou), and VI (Rio Negro)] were identified and found to be endemic in Florida, Mexico, and Central and South America (1, 2, 7–9). Of the thirteen viruses, VEEV-IAB and VEEV-IC are epizootic and have caused large outbreaks in equids and humans. The remaining eleven viruses are enzootic and can also cause severe disease in both equids and humans, however, they are not associated with large outbreaks.

VEEV-IAB has been recognized as an important pathogen in the Americas since the 1920s (1, 9, 10). Epizootics of VEEV-IAB were first recognized in Venezuela in 1936 and continued sporadically throughout the following four decades in South America, typically affecting tens to hundreds of thousands of people and resulting in hundreds of fatalities (1, 9–13). The most widespread VEEV-IAB outbreak occurred between 1969 and 1972, beginning near the Guatemala-El Salvador border on the Pacific coast and rapidly spreading through much of Central America before eventually reaching Texas (1, 9, 10). The outbreak was contained by the deployment of an experimental live-attenuated vaccine, TC-83, derived by serially passaging VEEV-IAB Trinidad donkey strain 83 times in guinea pig heart cells by the US military (9, 14).

VEEV is maintained in nature via a transmission cycle between *Culex (Melanoconion)* spp. and sylvatic rodents (9, 15–22). However, this cycle can spill over into equids and humans. The epizootic viruses of the VEE complex, subtypes IAB and IC, are more virulent in equids and humans, with mortality rates of 19% to 83% and <1%, respectively (1, 9). Children are more likely to develop fatal encephalitis and to suffer permanent neurological sequelae (1, 9). In addition, during the Cold War, VEEV-IAB was developed as a threat agent for aerosol dissemination by both the United States of America (USA) and the Union of Soviet Socialist Republics (USSR) (23, 24). These traits have led to the assignment of VEEV-IAB and VEEV-IC to the NIAID Category B pathogens and are categorized as select agents. Currently, no licensed therapeutics or vaccines are available to treat or prevent VEEV infection, and the US population remains vulnerable to natural disease outbreaks and bioterror events.

Limited data are available on VEEV infection in humans. Most available data derive from accidental exposures in laboratory settings via aerosol or percutaneous routes (25). The infectious dose of VEEV in these human cases remains unknown. Nonetheless, reported symptoms typically include acute febrile illness, usually with fever, chills, headache, back pain, malaise, myalgia, anorexia, nausea, and mild encephalitis. The lack of a detailed disease course in humans is a critical impediment in understanding VEEV pathogenesis as well as countermeasure development. The cynomolgus macaque model following VEEV-IAB infection has been used to recapitulate aspects of human disease and to provide insight into viral pathogenesis (26–40). Cynomolgus macaques do not exhibit overt disease via peripheral routes, however, infection with aerosol route alters body temperature, blood chemistry, and hematology (27–29, 32, 33, 35, 38, 41). However, the data in cynomolgus macaques are limited and the model requires additional development to understand VEEV-IAB pathogenesis. In this proof-of-concept study we investigated the disease course of VEEV-IAB Trinidad donkey (TrD) strain in the cynomolgus macaque model by utilizing state-of-the-art telemetry to measure disease signs including temperature, activity, respiration, heart rate, blood pressure, electrocardiography (ECG), and electroencephalography (EEG) following an aerosol infection.

## RESULTS

### VEEV-IAB study design

Four cynomolgus macaques (one male and three females) were implanted with telemetry devices to monitor physiological parameters including temperature, respiration, heart rate, blood pressure, activity, ECG, and EEG simultaneously and continuously in a 33-day study (Fig. 1). The baseline for each parameter was established in 48 half-hour intervals per day by averaging raw data points from five pre-infection daytime and nighttime cycles. Day and night times were defined as 6 am to 6 pm and 6 pm to 6 am, respectively. The baseline for each parameter is plotted as a grey line for each individual NHP in the figures. All post-infection comparisons were time-matched to respective day or nighttime pre-infection values. In addition, NHPs were also monitored for disease signs and alteration in behavior pre- and post-infection. Following establishment of baseline, NHPs were infected via the aerosol route with VEEV-IAB TrD strain at a target dose of 6.0 log_10_ PFU (Fig. 1A). NHPs received virus doses ranging from ∼6.2 to 6.5 log_10_ PFU (Fig. 1B).

**Figure 1.**
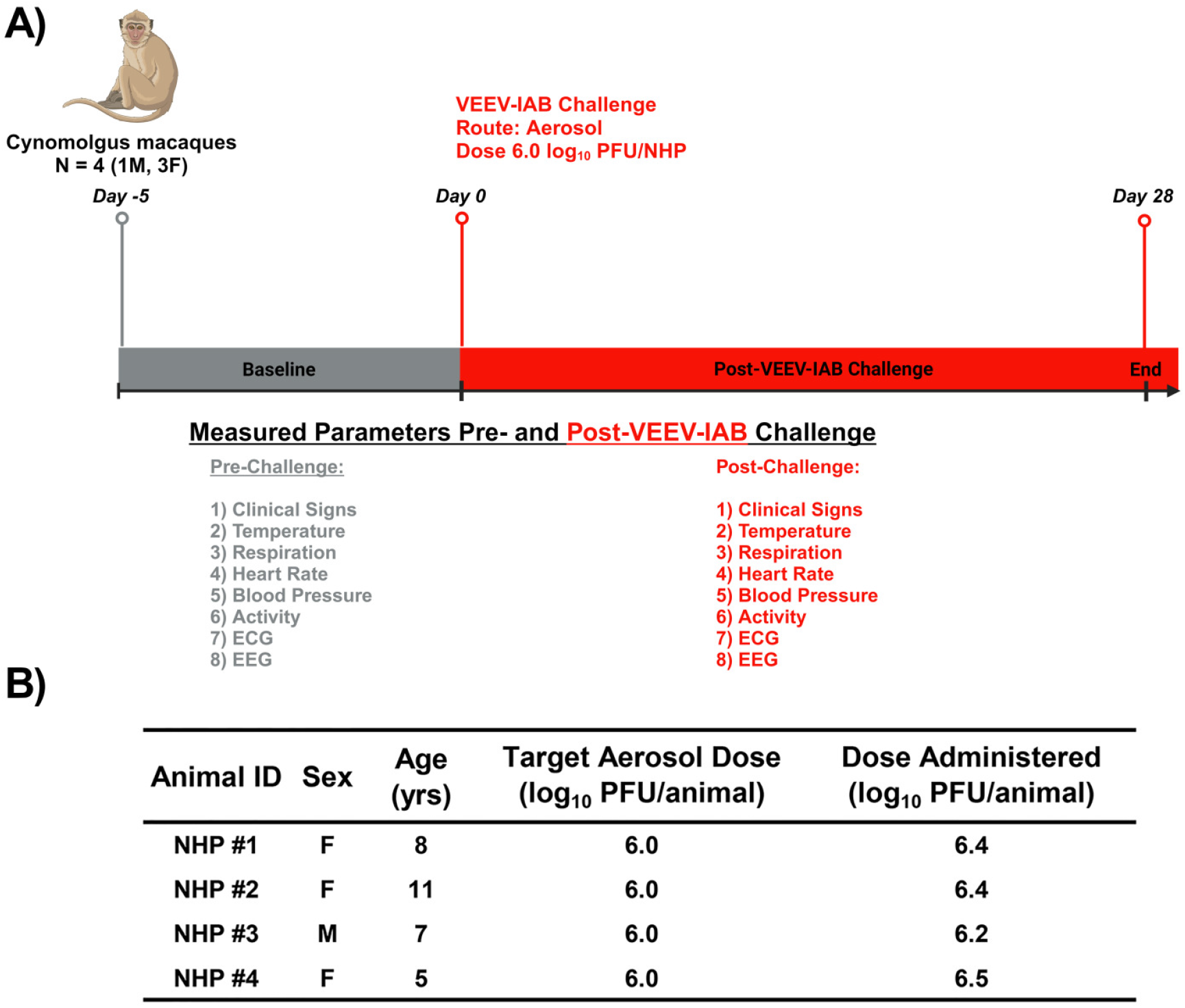
Experimental design of the cynomolgus macaque study (A). NHP weights, ages, target and experimental dose (B).

### Disease Scores, Alteration in Behavior, and Activity

Following infection, NHPs were observed for signs of disease, and each animal was assigned a score based on alterations in temperature, responsiveness, and neurological changes. All NHPs exhibited signs of disease by 2 dpi (Supp. Fig. 1). The peak score was ∼2 to 5 at 6 dpi and all NHPs returned to baseline by 7 to 8 dpi. No overt neurological signs were observed in any of the NHPs post-VEEV-IAB infection.

The NHPs were continuously monitored remotely with limited disruptions from human activity. This provided a rare opportunity to study the impact of VEEV-IAB infection on animal behavior. Baseline behavior was established for each animal over five days prior to infection during both daytime and nighttime cycles. Alteration in animal behavior was assessed by observing three parameters: sleep, activity, and food/fluid consumption. Disruption of circadian rhythm was detected as early as ∼1 dpi in NHP #1 with increased activity at nighttime and a corresponding decrease in sleep (Supp. Table 1). All NHPs exhibited a decrease in sleep by the 2^nd^ and 3^rd^ nighttime periods, which persisted throughout 4 to 9 dpi. Reduced sleep was accompanied by decreased daytime activity in NHPs #1, 2 and 3. The reduced daytime activity in NHP #2 at 8 dpi was due to several prolonged sleep periods during the day. Additionally, NHPs #1-3 had a marked reduction in fluid consumption between 5 to 9 dpi. By 11 dpi, all NHPs returned to baseline sleep, activity, and fluid consumption.

**Table 1.** Summary of fever data in NHPs infected with VEEV-IAB via aerosol route. Fever hours are calculated as the sum of the significant temperature elevations. ΔT_max_ = maximum change in temperature.

| Animal # | Onset of Fever<br>(hpi) | Fever Duration<br>(hrs) | Fever<br>Hours | $\Delta T_{\max}$<br>(°C) | Peak Temperature<br>(°C) |
| --- | --- | --- | --- | --- | --- |
| NHP #1 | 18 | 97.5 | 277 | 3.9 | 40.8 |
| NHP #2 | 31 | 88.5 | 261 | 3.8 | 40.6 |
| NHP #3 | 24 | 138 | 397 | 4 | 40.7 |
| NHP #4 | 48 | 45.5 | 181 | 3.1 | 40.1 |

The NHP activity was also measured and quantitated via telemetry by recording accelerations in X, Y, and Z dimensions, which were combined to generate an index with arbitrary units proportional to the three-dimensional movements. Baseline activity ranged from ∼329 to 748 units during the daytime and ∼320 to 393 units at night (Supp. Fig. 2). Daytime activity decreased in all NHPs from 1 to 10 dpi, with peak reductions of ∼29–55%. In contrast, the nighttime activity increased between 2 to 9 dpi with peak values of ∼+14 to +34%. The increased nighttime activity was more pronounced for NHPs #2 and #4. NHP #2 exhibited increased activity at 5 dpi throughout the entire nighttime period with peak increase of ∼+31%. NHP #4 exhibited increased nighttime activity throughout 1 to 9 dpi with peak increase of ∼+29%. By 11 dpi, activity levels in all NHPs had returned to baseline.

**Figure 2.**
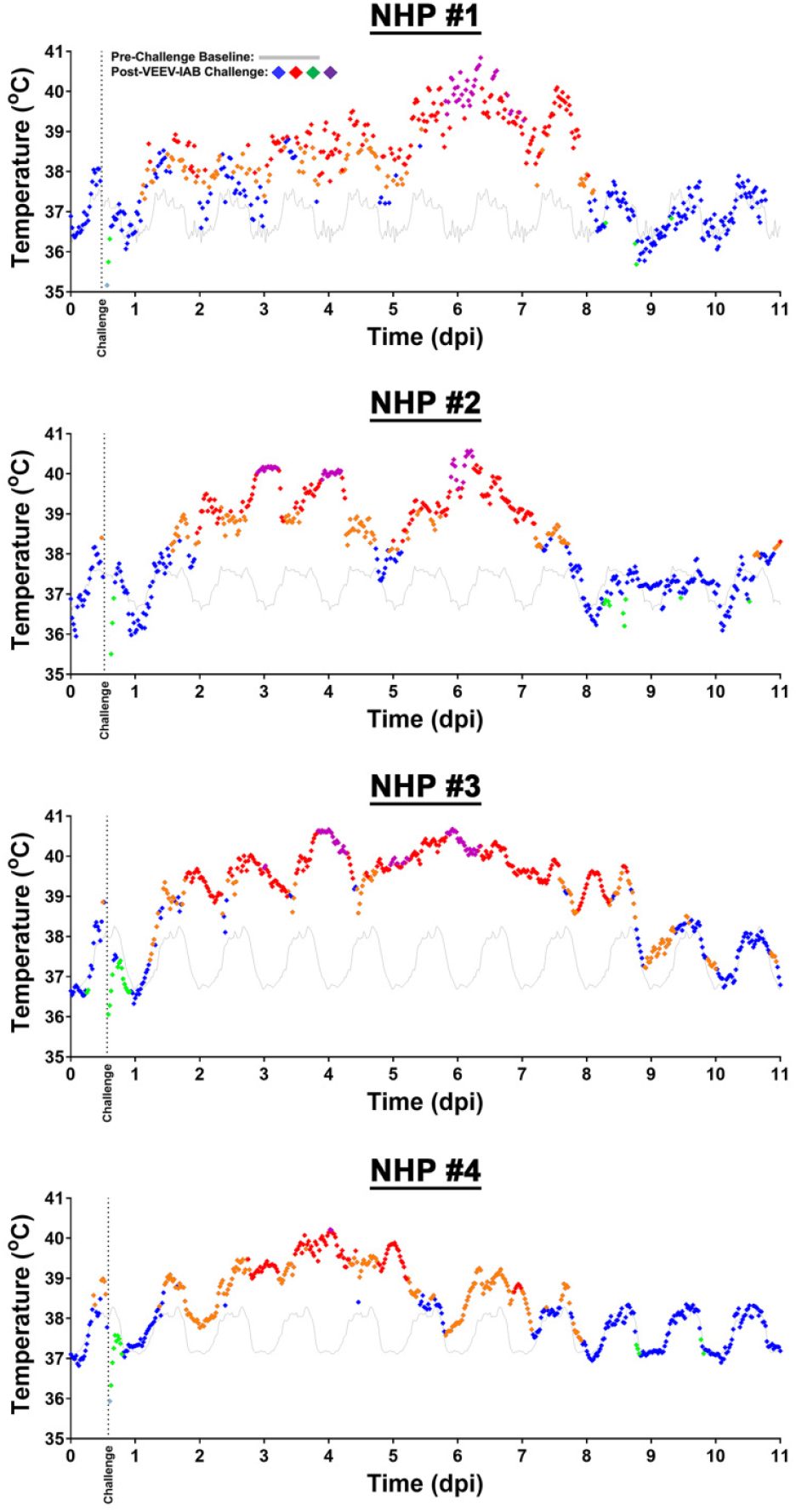
Body temperature of NHPs pre- and post-VEEV-IAB challenge. Data analysis is shown in 0.5-hr intervals. Pre-challenge baseline temperature is shown in grey. Values within ≤3 standard deviations (SD) are indicated with (♦), >3 SD above baseline are indicated with (♦), and <3 SD below baseline are indicated with (♦). Hyperpyrexia is indicated by (♦).

### Temperature and Respiration

Following infection, three of the four NHPs exhibited a rapid temperature increase within 1 dpi (Fig. 2 and Table 1). The onset of fever occurred within ∼1 to 2 dpi in all NHPs, with fever duration of ∼46 to 138 and total fever hours of ∼181 to 397. The ΔT_max_ was ∼3.1 to 4° C, with peak temperature of ∼40.1 to 40.8° C (Table 1). The temperature returned to baseline between 8 to 10 dpi in all NHPs.

The baseline daytime and nighttime respiration rate ranged from ∼19 to 32 and ∼16 and 24 bpm, respectively (Fig. 3). Respiration rate increased during both day and nighttime periods between 2 to 10 dpi. Peak daytime increases ranged from ∼+30 to +63%, while nighttime increases were more pronounced with values ranging from ∼+45% to +91%. NHP #1 exhibited a reduction in daytime respiration between 8 and 9 dpi of ∼-25%. NHPs 2 and #3 consistently exhibited elevated respiration rates and did not return to baseline values by 11 dpi, whereas NHPs #1 and #4 returned to near baseline.

**Figure 3.**
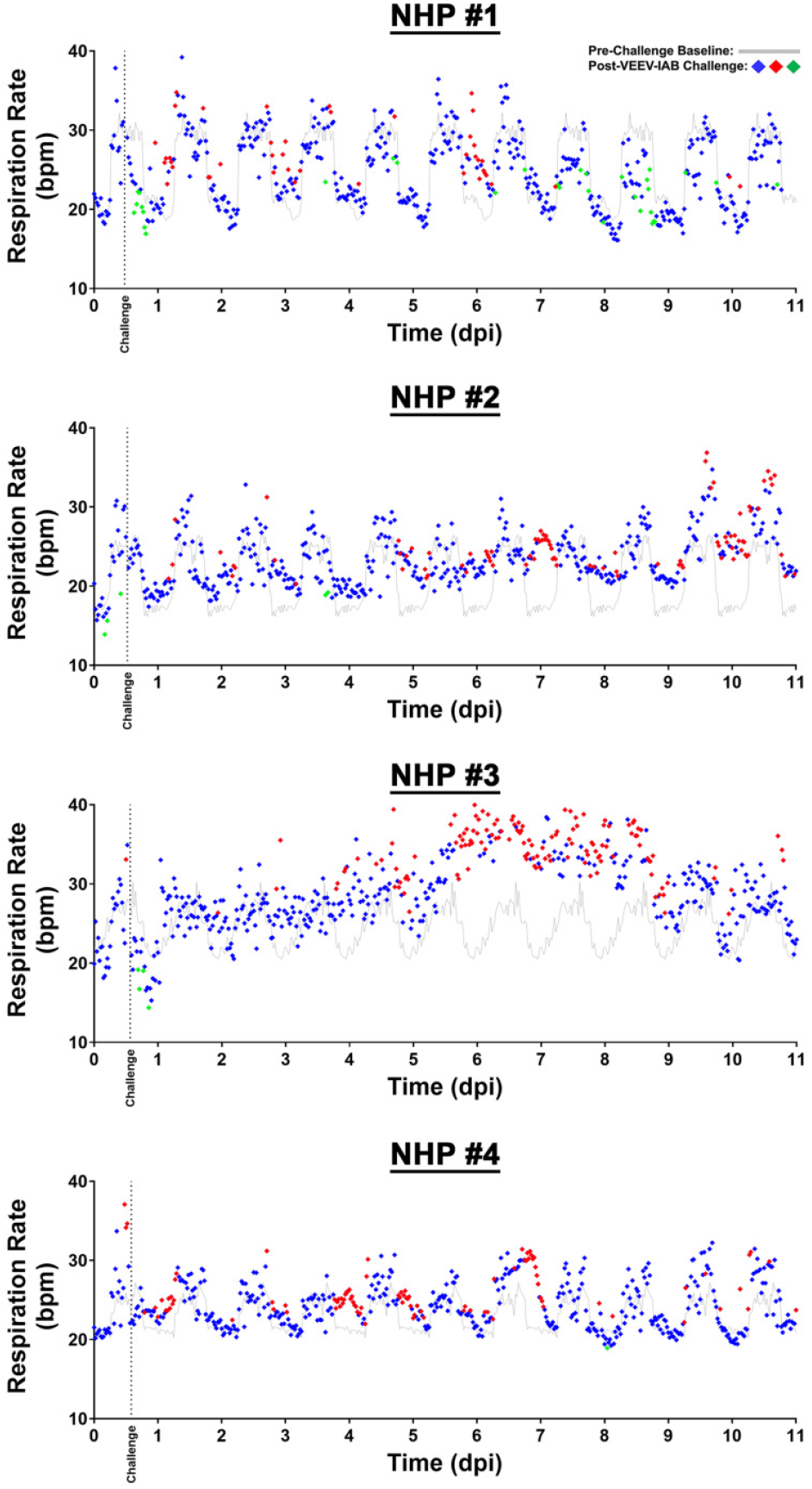
Respiration rate of NHPs pre- and post-VEEV-IAB challenge. Data analysis is shown in 0.5-hr intervals. Pre-challenge baseline respiration rate is shown in grey. Values within ≤3 standard deviations (SD) are indicated with (♦), >3 SD above baseline are indicated with (♦), and <3 SD below baseline are indicated with (♦). bpm = breaths per minute. *p*-values are indicated via (♦) = *p <0.003* and (♦) = *p <0.02*.

### Heart Rate, Blood Pressure, and ECG

The baseline day and nighttime heart rates ranged from ∼76 to 175 and ∼63 to 153 bpm (Fig. 4). Following infection, three distinct response patterns emerged. NHP #1 exhibited a sustained daytime decrease (∼-27 to -40%) with a concurrent increase at nighttime rate (∼+16 to +60%). NHP#2 maintained a near normal daytime rate with an increase in nighttime rate (∼+13 to +56%) throughout 11 dpi. In contrast, NHPs #3 and #4 exhibited sustained increases in both day and nighttime heart rates. NHP #3 exhibited the greatest elevations, with daytime increases of ∼+26 to +149% and nighttime increases of ∼+41 to +191%. NHP #4 showed more moderate increases, with daytime and nighttime peaks of ∼+18 to +57%. None of the NHPs returned to baseline values by 11 dpi.

**Figure 4.**
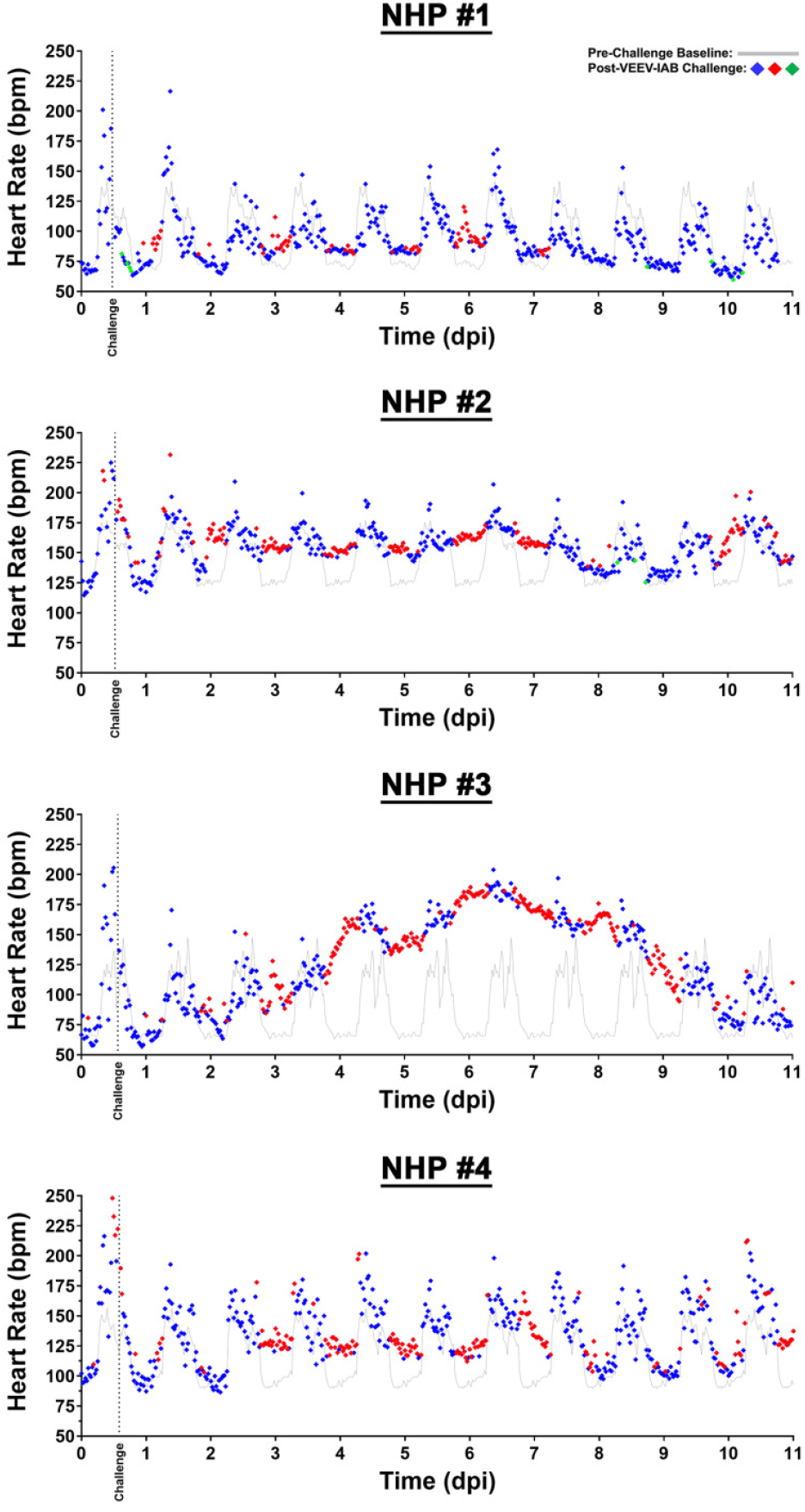
Heart rate of NHPs pre- and post-VEEV-IAB challenge. Data analysis is shown in 0.5-hr intervals. Pre-challenge baseline respiration rate is shown in grey. Values within ≤3 standard deviations (SD) are indicated with (♦), >3 SD above baseline are indicated with (♦), and <3 SD below baseline are indicated with (♦). bpm = beats per minute. *p*-values are indicated via (♦) = *p <0.002* and (♦) = *p <0.01*.

Systolic and diastolic blood pressure were measured in NHPs #1, 2, and 4 (Supp. Fig. 3 and 4). All NHPs exhibited intermittent alterations in blood pressure throughout 11 dpi, however, all NHPs exhibited elevated blood pressure during nighttime. Daytime systolic blood pressure varied, with increases of ∼+6 to +13% and decreases of ∼-5 to -18% (Supp. Fig. 3). During nighttime, systolic blood pressure remained elevated by ∼+11 to +39%. Diastolic blood pressure exhibited a similar pattern to systolic blood pressure, with daytime blood pressure increased or decreased of ∼+6 to +15% and ∼-6 to -20%, respectively (Supp. Fig. 4). The increase in nighttime blood pressure ranged from ∼+7 to +39% (Supp. Fig. 4). Neither systolic nor diastolic blood pressure returned to baseline in all NHPs at 11 dpi.

Cardiac electrical activity was assessed via ECG to measure RR (single heartbeat), PR (atrial contraction), QRS duration (ventricle contraction), and QTc Bazett (time between ventricle relaxation and contraction) (Supp. Fig. 5 and 6). The sustained elevation in heart rate was associated with reduction in all intervals (Supp. Fig. 5 and 6). However, in both NHPs #3 and #4 ECG abnormalities were observed. Both NHPs exhibited increased PR duration between 8 and 10 dpi, while NHP #3 had a marked increase in QRS duration between 3 to 11 dpi and QTc Bazett between 7 to 10 dpi.

**Figure 5.**
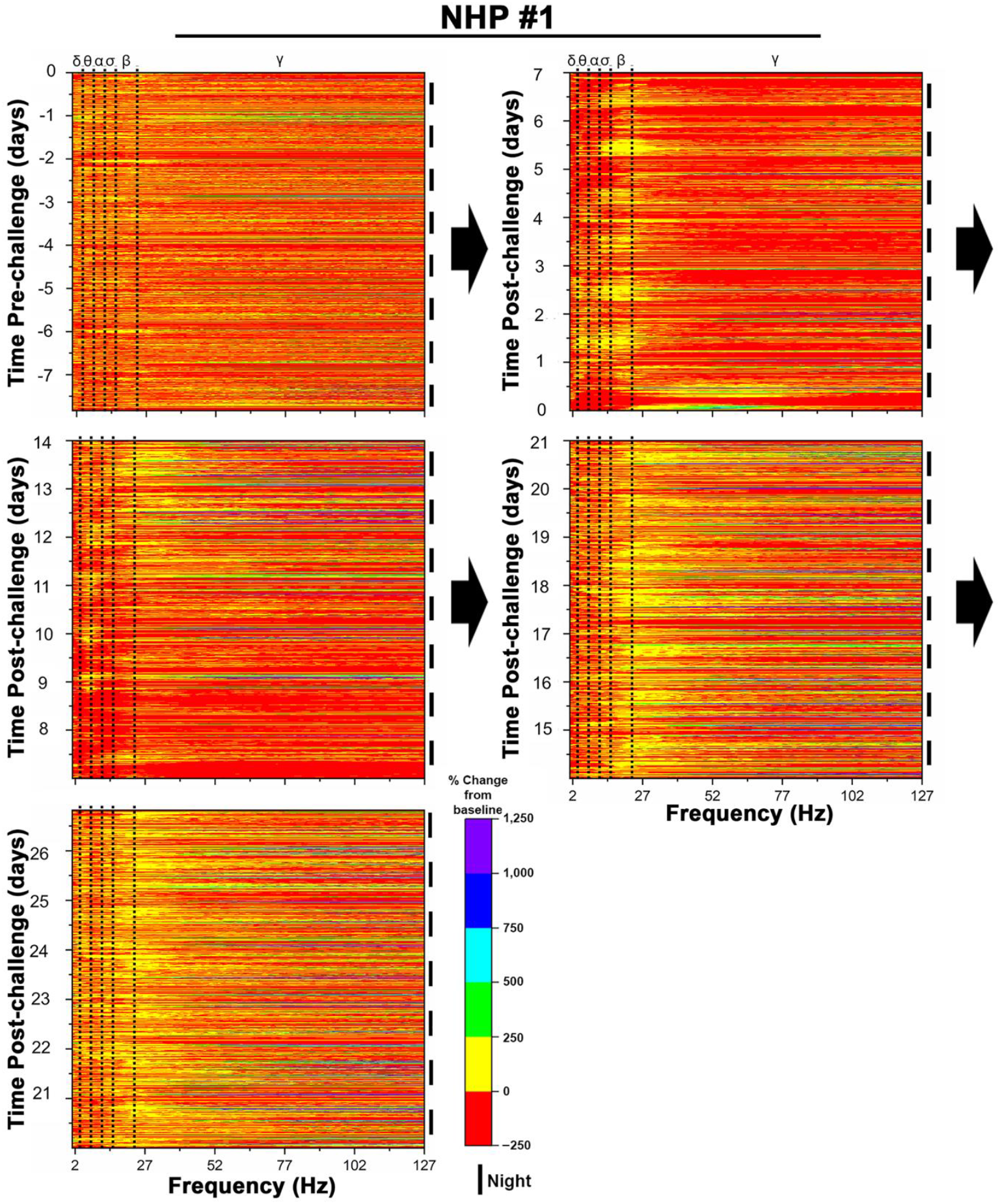
Pre- and post-VEEV-IAB challenge quantitative electroencephalography (qEEG) heat maps of NHP #1. The top and bottom x- axes display brain waves [delta (δ), theta (θ), alpha (α), sigma (σ), and gamma (γ)] and frequency in hertz (Hz), respectively. The left and right y-axes display time (days) and 12-hr day/nighttime intervals, respectively. Pre-challenge baseline (top left) and post-challenge heat maps are shown via arrows. 12-hr nighttime is indicated by (I) and daytime by a gap.

**Figure 6.**
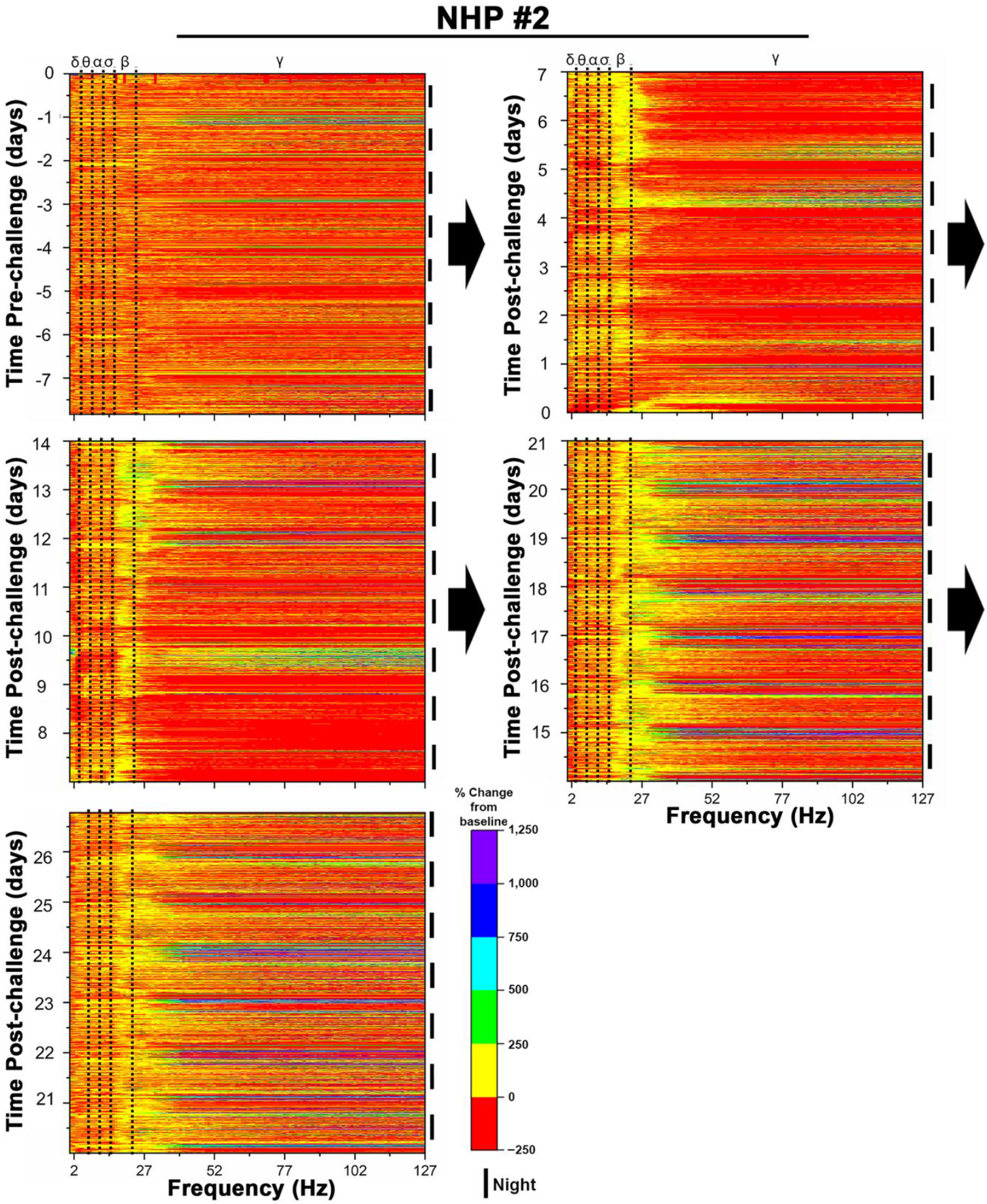
Pre- and post-VEEV-IAB challenge quantitative electroencephalography (qEEG) heat maps of NHP #2. The top and bottom x- axes display brain waves [delta (δ), theta (θ), alpha (α), sigma (σ), and gamma (γ)] and frequency in hertz (Hz), respectively. The left and right y-axes display time (days) and 12-hr day/nighttime intervals, respectively. Pre-challenge baseline (top left) and post-challenge heat maps are shown via arrows. 12-hr nighttime is indicated by (I) and daytime by a gap.

### Quantitative EEG (qEEG)

The qEEG data for each NHP pre- and post-infection is shown as heat maps (Figs. 5-8). Pre-infection baselines were generated from eight nighttime and seven daytime 12-hr intervals. Absolute powers of frequency bands delta (δ) [0.5-4 Hz], theta (θ) [4-8 Hz], alpha (α) [8-12 Hz], sigma (σ) [12- 16-24 Hz], beta (β) [16-24 Hz] and gamma (γ) [24-50 Hz] were calculated using NeuroScore software. In all four NHPs, baseline heat maps displayed similar patterns, with most values between ∼-50 to +50%, and few exceeding 100%. Increases in gamma waves were observed during the daytime typically coinciding with anticipated feeding periods and/or other scheduled activities in all four NHPs and were attributable to electromyographic artifacts.

Immediately following the aerosol challenge [∼4 to 10 hours post-infection (hpi)], the uniform distribution observed during the baseline period for all NHPs was replaced with an increase in lower frequency intensities (i.e., <27 Hz). These early changes were associated with administration of sedative drugs (i.e., Ketamine, Telazol) at the time of aerosol infection. Nonetheless, shortly after this period of sedative-induced reduction in individual frequencies, distinct patterns could be rapidly observed across all NHPs.

Following recovery from anesthesia at 12 hpi, NHP #1 exhibited prolonged periods of increase (∼+250%) in both sigma and beta waves at nighttime throughout the 1^st^ week post-infection (Fig. 5). During this period, there was a marked decrease in delta, theta, alpha, and gamma waves between 3 and 7 dpi in both day and nighttime periods. During week two-post infection, all waves declined by up to ∼-250% from baseline values between 7 and 9 dpi. At 10 to 14 dpi, daytime increases of ∼+250% in delta and theta waves were observed, with peak increases of ∼+500%. Concurrently, a marked and sustained increase of ∼+250% in beta and low-frequency gamma (up to 40 Hz) waves were observed. Week 3 post-infection was characterized by sustained increases of ∼+250% in beta and low-frequency gamma (up to 40 Hz). High frequency gamma waves (> 40 Hz) were punctuated with sporadic increases of >+1,000% relative to baseline values. This pattern persisted into week 4 post-infection, with sustained increases of ∼+250% in beta and low-frequency gamma waves accompanied by variable increases of up to ∼+1,000% in high frequency gamma waves (> 40 Hz). At 27 dpi NHP #1 brain wave activity, particularly the beta and gamma waves, did not return to baseline.

NHP #2 exhibited nighttime increases of up to ∼+250% in delta, theta, alpha, sigma, and beta waves throughout week one post-infection (Fig. 6). The increase was more pronounced and prolonged in the beta power band between 4 to 6 dpi. Gamma wave activity increased by ∼+250% to +750% at nighttime between 4 and 5 dpi. During week two post-infection, low-frequency (<27 Hz) waves exhibited prolonged and sustained increases of up to ∼+500% in beta waves. Gamma waves displayed a biphasic trend with decreases of up to ∼- 250% between 7 and 9 dpi, followed by protracted increases of up to ∼+4,800% most notably at nighttime on 9 dpi. During third- and fourth-weeks post-infection, the brain activity exhibited a similar pattern to week 2 with sustained increases of ∼+250 to +1,000% in beta and gamma waves throughout both weeks. At 27 dpi NHP #2 brain wave activity, particularly the beta and gamma waves, did not return to baseline.

NHP #3 exhibited a marked increase of up to ∼+250% to +500% in delta, theta, alpha, sigma, and beta waves throughout the first week post-infection, with increases in beta waves more pronounced during nighttime periods (Fig. 7). Low-frequency gamma waves (<40 Hz) exhibited an increase of ∼+250% at nighttime throughout 7 dpi, whereas high frequency gamma waves (>40 Hz) declined between 4 and 7 dpi. Throughout week two post-infection, the gamma waves exhibited a prolonged and intense increase of up to ∼+750% to +4,000% at nighttime followed by a decrease during daytime of up to ∼-250%. Similar to week one, delta, theta, alpha, and sigma waves continued to exhibit sustained increase of ∼+250% to +500% throughout 10 dpi. Beta waves exhibited an intense increase at nighttime of ∼+250% between 7 to 10 dpi. This was followed by a gradual and sustained decline in beta waves throughout the remainder of week two post-infection. Week three post-infection was marked by intermittent fluctuations across all bands, however, beta and gamma waves exhibited pronounced increases of ∼+250 to +500% at nighttime as well as at night-to-day transitions. A similar pattern persisted into week four post-infection, delta, theta, alpha, and sigma waves exhibiting intermittent increases of up to ∼+500%. Intense increases in beta waves continued throughout the nighttime periods. Lastly, gamma waves exhibited increases of ∼+250% to +1,000%. At 27 dpi NHP #3 brain wave activity, particularly the beta and gamma waves, did not return to baseline.

**Figure 7.**
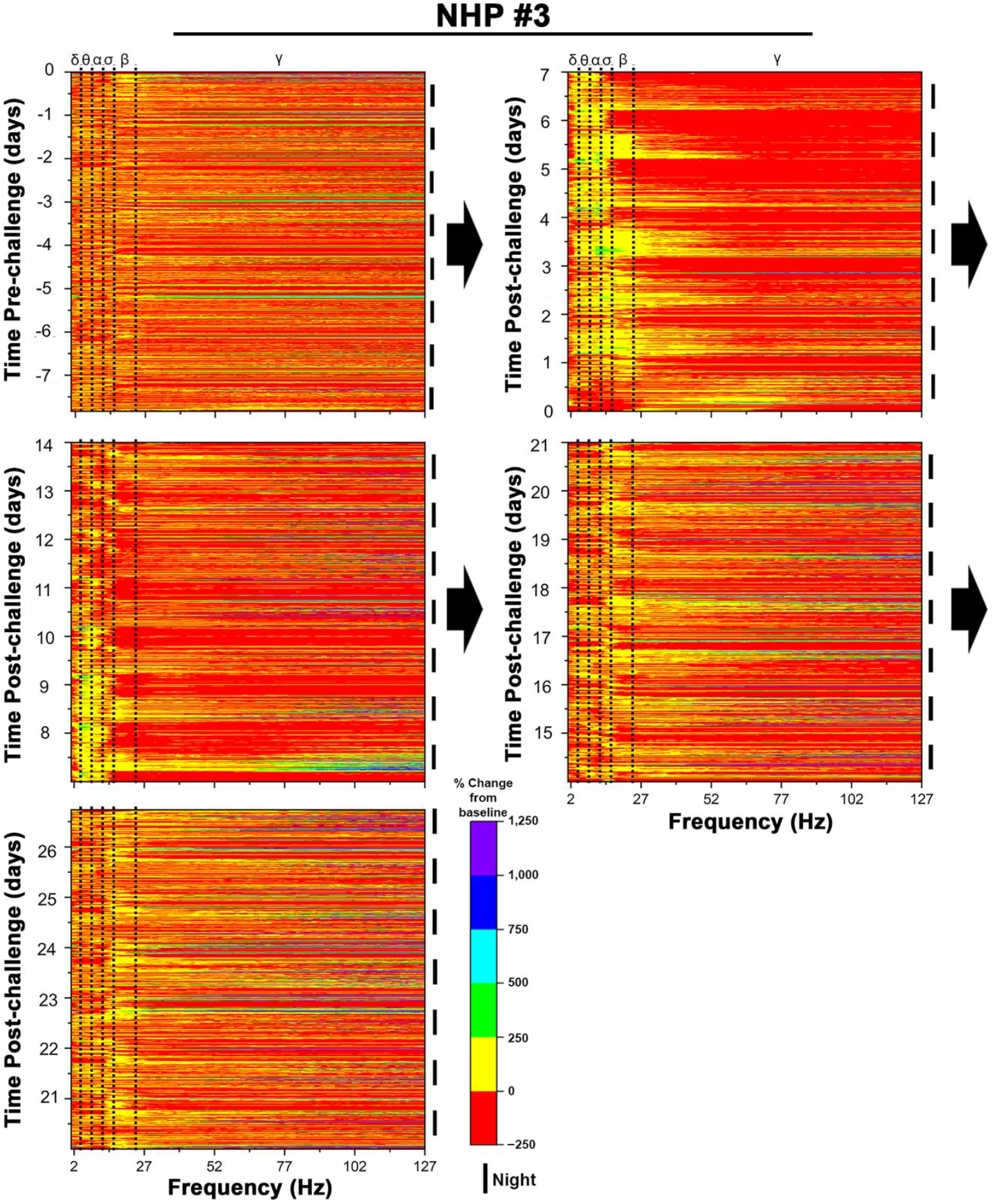
Pre- and post-VEEV-IAB challenge quantitative electroencephalography (qEEG) heat maps of NHP #3. The top and bottom x- axes display brain waves [delta (δ), theta (θ), alpha (α), sigma (σ), and gamma (γ)] and frequency in hertz (Hz), respectively. The left and right y-axes display time (days) and 12-hr day/nighttime intervals, respectively. Pre-challenge baseline (top left) and post-challenge heat maps are shown via arrows. 12-hr nighttime is indicated by (I) and daytime by a gap.

**Figure 8.**
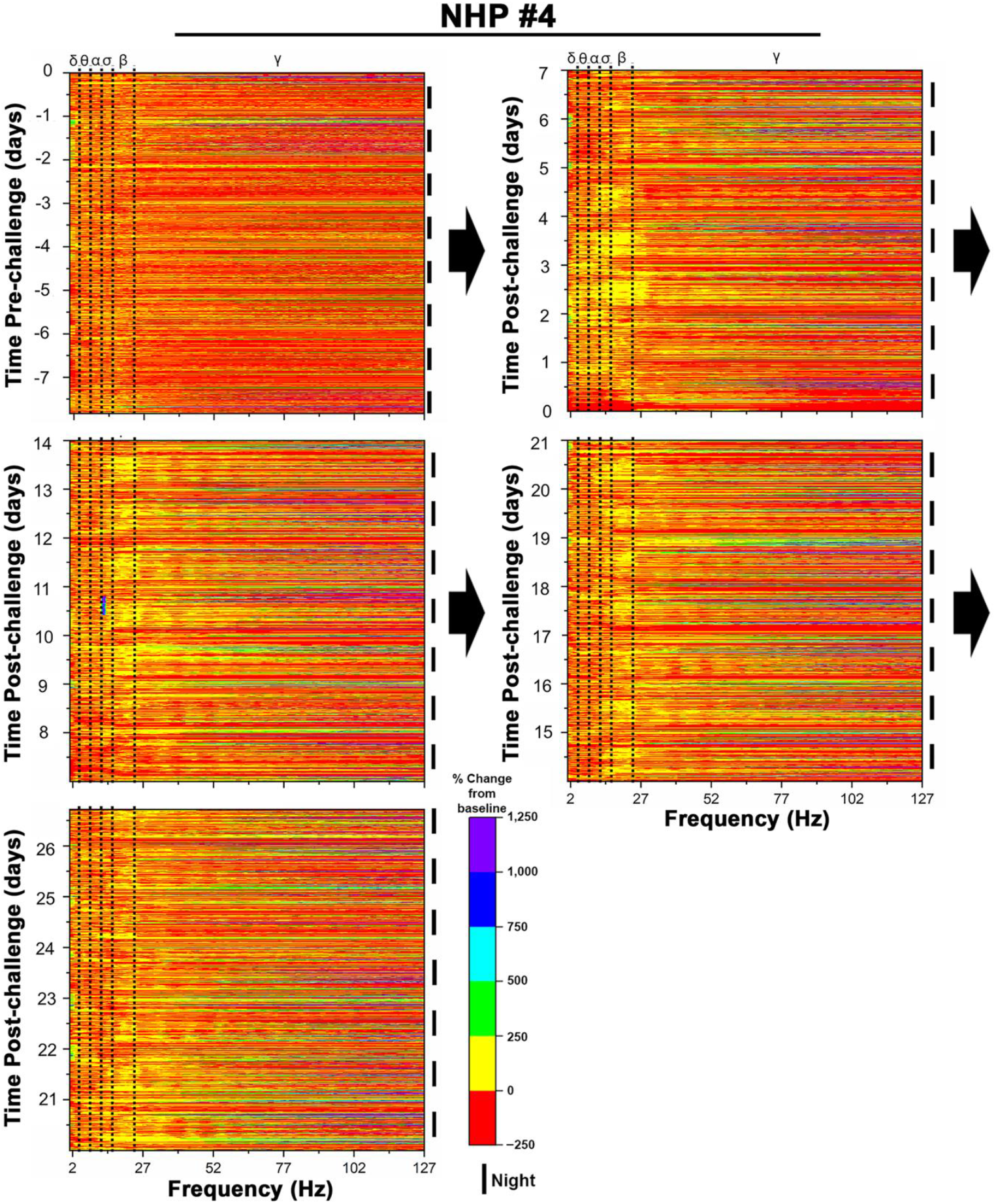
Pre- and post-VEEV-IAB challenge quantitative electroencephalography (qEEG) heat maps of NHP #4. The top and bottom x- axes display brain waves [delta (δ), theta (θ), alpha (α), sigma (σ), and gamma (γ)] and frequency in hertz (Hz), respectively. The left and right y-axes display time (days) and 12-hr day/nighttime intervals, respectively. Pre-challenge baseline (top left) and post-challenge heat maps are shown via arrows. 12-hr nighttime is indicated by (I) and daytime by a gap.

NHP #4 exhibited broad increases of ∼+250% to +500% in delta, theta, alpha, sigma, and beta waves between 1 to 4 dpi (Fig. 8). Delta waves showed intermittent daytime increase of ∼+500% between 5 and 6 dpi. Concurrent with these alterations, the gamma waves exhibited highly recurrent increases of up to ∼+1,000% throughout the first week post-infection. Throughout week two and three post-infection, increases in beta and gamma waves remained elevated. Beta waves increased up to ∼+250%, particularly at nighttime between 9 to 20 dpi. While gamma wave activity rose by ∼+750% to +1,000% relative to baseline value throughout the two-week period. During week four post-infection, beta activity remained elevated but began to decline gradually after 23 dpi. Similarly, gamma waves also remained elevated, with increases of up to ∼+750%. At 27 dpi NHP #4 brain wave activity, particularly the beta and gamma waves, did not return to baseline.

### Pathology

Tissues from all NHPs were harvested at the end of the study at 28 dpi. Brain and spinal cord tissue samples were collected, and various regions were sectioned to investigate VEEV-IAB induced pathology. The brain regions included frontal cortex, amygdala, hippocampus, hypothalamus, thalamus, corpus striatum, mesencephalon, medulla oblongata, and cerebellum. Spinal cord regions included cervical, thoracic, and lumbar samples. Minimal or no macroscopic lesions were observed in all NHP tissues (Figs. 9 and 10). In addition, *in situ* hybridization (ISH) and immunohistochemistry (IHC) were unable to detect the presence of viral RNA or proteins in most brain regions, respectively. However, VEEV-IAB RNA was detected in rare sections of the brain (Fig. 11).

**Figure 9.**
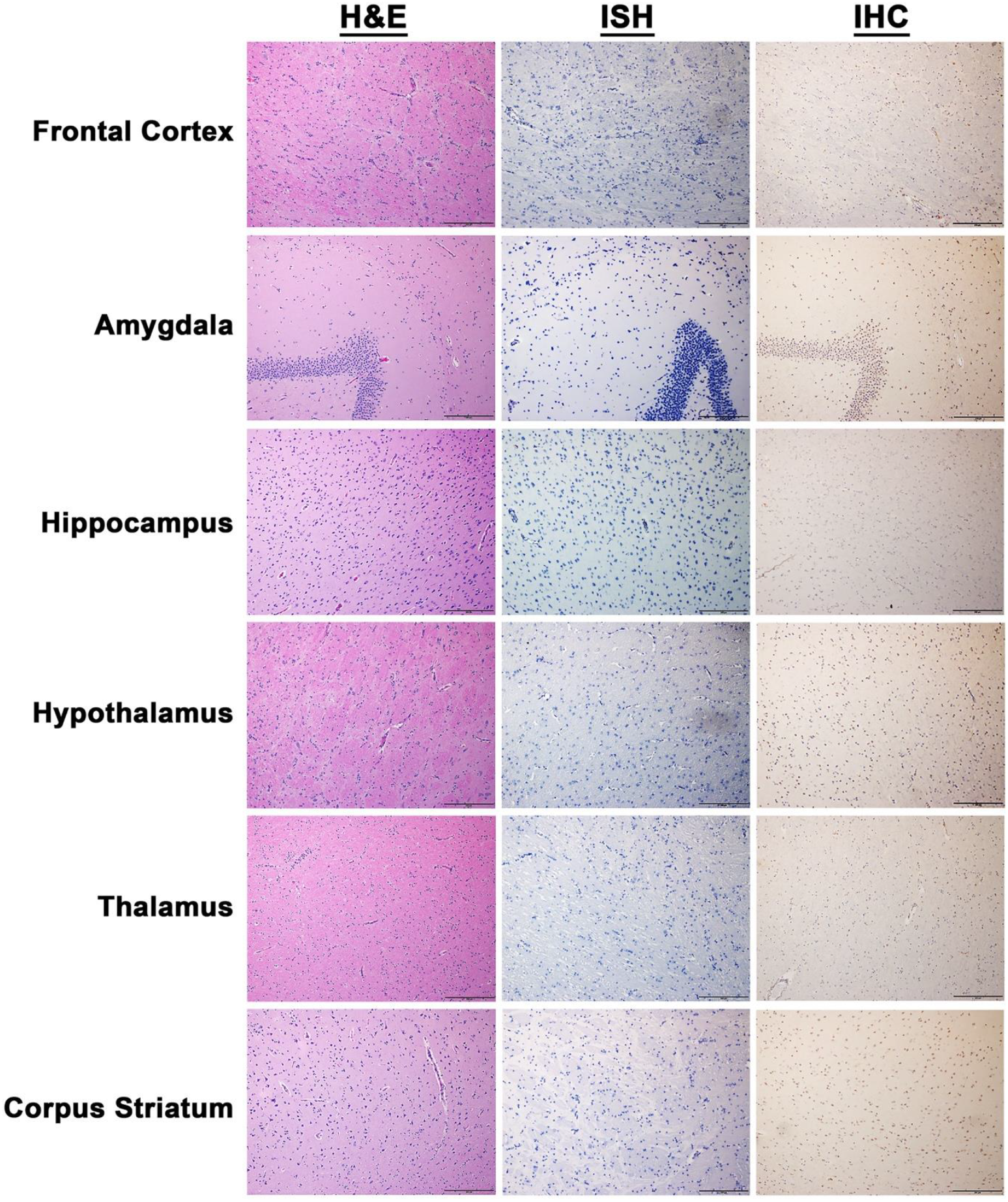
Pathology in the brain regions from the NHPs at 28 dpi. The tissues were collected at the time of euthanasia. Hematoxylin and eosin (H&E) staining was performed to visualize histopathology. The presence of VEEV-IAB RNA and proteins was visualized via *in situ* hybridization (ISH) and immunohistochemistry (IHC), respectively. Representative pictures are shown.

**Figure 10.**
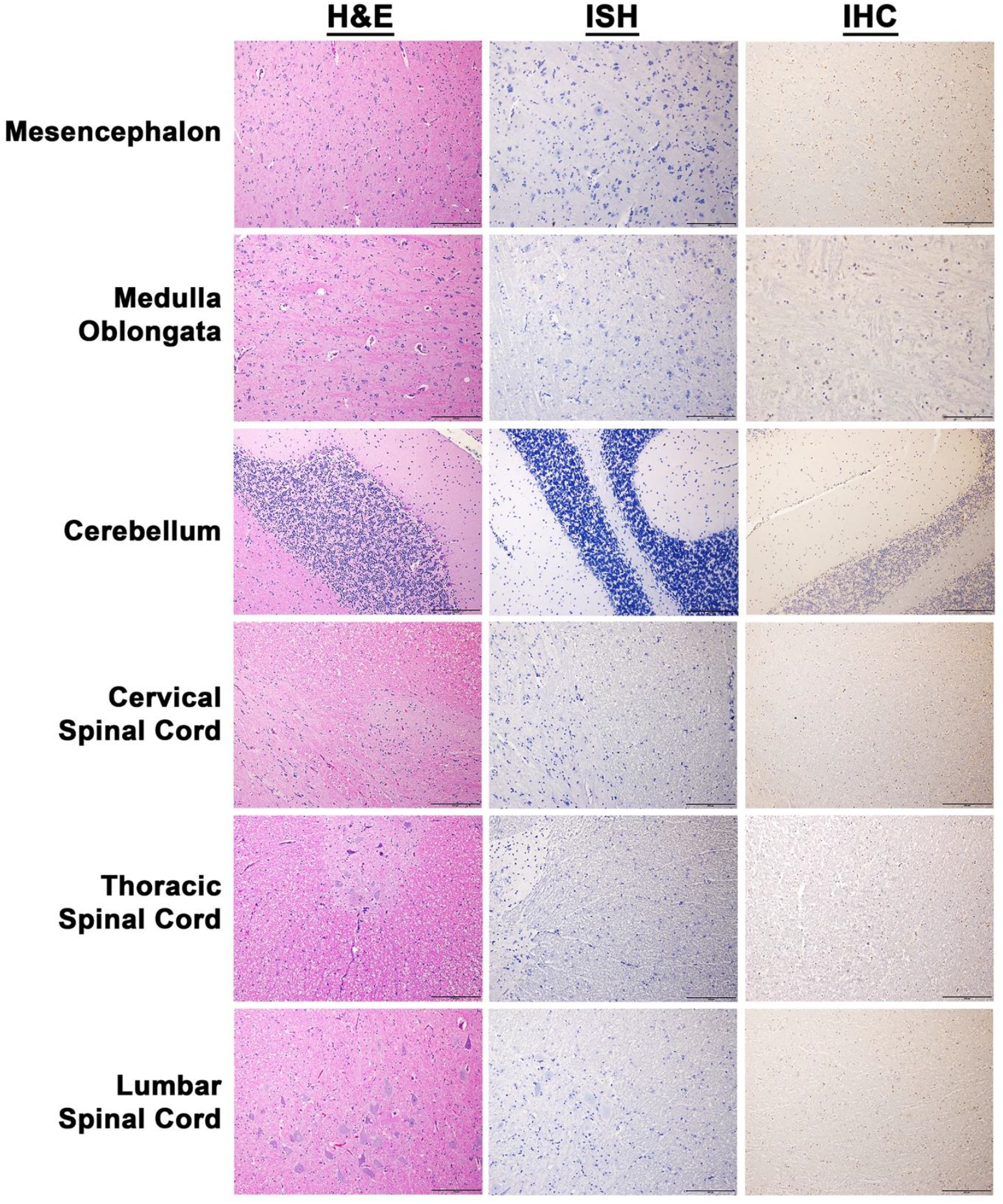
Pathology in the brain and spinal cord regions from the NHPs at 28 dpi. The tissues were collected at the time of euthanasia. Hematoxylin and eosin (H&E) staining was performed to visualize histopathology. The presence of VEEV-IAB RNA and proteins was visualized via *in situ* hybridization (ISH) and immunohistochemistry (IHC), respectively. Representative pictures are shown.

**Figure 11.**
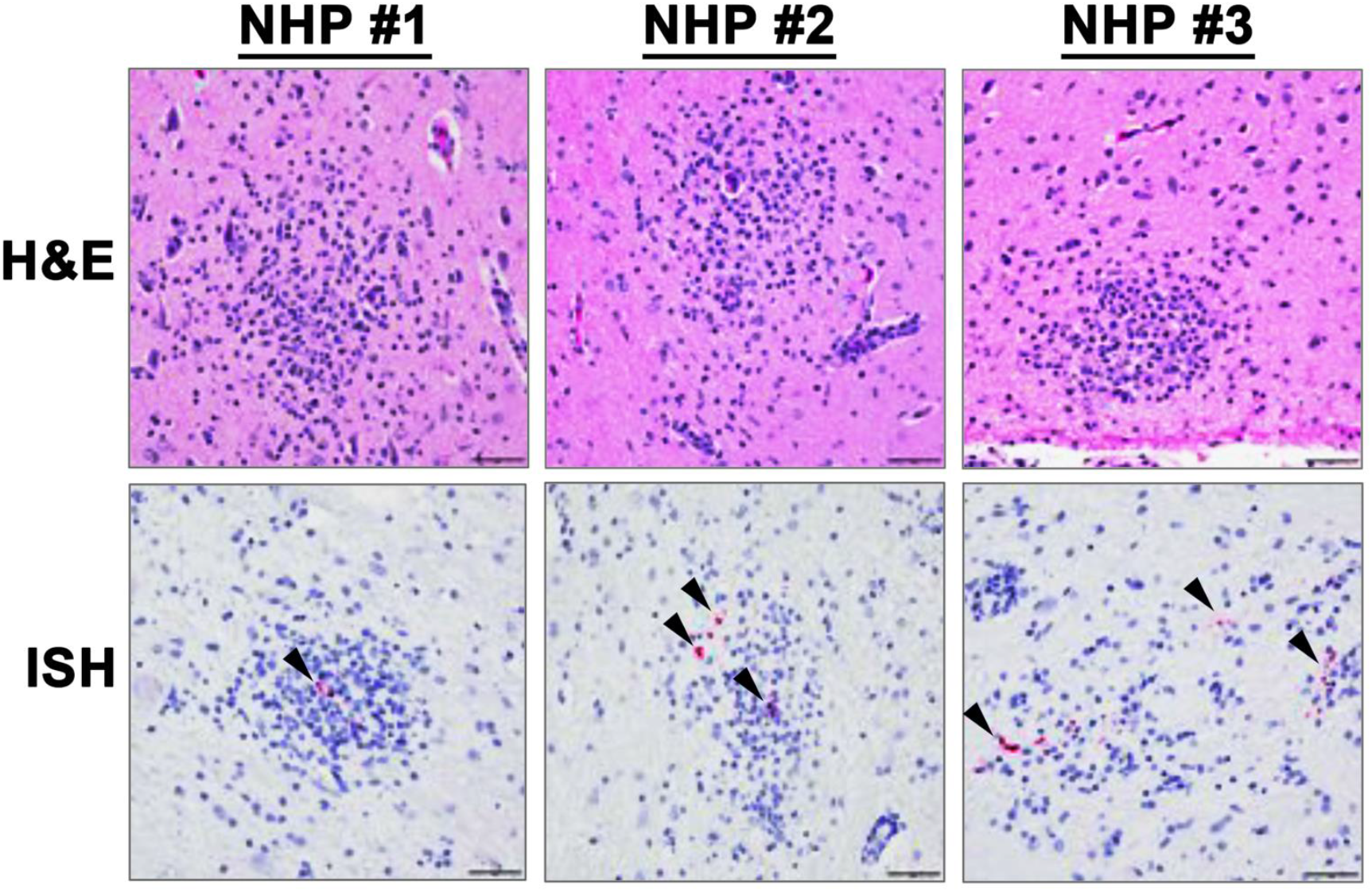
*in situ* hybridization (ISH) to detect VEEV-IAB RNA in brain tissues of three NHPs at 28 dpi. The tissues were collected at the time of euthanasia.

### Neutralizing antibody response

The neutralizing antibody responses were assessed via PRNT_80_ at days -7, 0, and terminal time points (Supp. Table 2). None of the NHPs had detectable neutralizing antibody titers prior to or at the time of infection and were assigned the limit of detection of the assay (<1:20). By 28 dpi, all NHPs developed robust neutralizing antibody responses, with PRNT₈₀ titers exceeding 1:5,120 (Supp. Table 2).

## DISCUSSION

State-of-the-art telemetry offers considerable advantages for the development of next-generation animal models, particularly for studies involving Risk Group 3 and 4 agents. First, telemetry permits continuous and simultaneous monitoring of multiple critical physiological parameters for up to at least six weeks. Second, it generates high-quality, high-resolution data, with up to 1,000 data points collected per second for individual parameters (42). Third, telemetry decreases or eliminates the need for frequent animal handling, thereby reducing animal stress and pain. It also minimizes or eliminates the need for study personnel to enter high-containment facilities, thereby enhancing safety while reducing time and operational costs. Fourth, it can reduce the number of animals required for studies. Physiological parameters are influenced by factors such as sex, age, and genetics, and are therefore inherently variable between animals (43–48). This inter-animal variability increases the number of animals required to achieve sufficient statistical power in animal studies. Since telemetry provides high-resolution longitudinal data, each animal can serve as its own control through pre- and post-treatment comparisons, thereby reducing animal numbers while increasing statistical power (Supp. Table 3). These latter advantages are aligned with the principles of the 3Rs (Replacement, Reduction, and Refinement), which aim to improve the ethics and welfare of animals used in scientific research.

While telemetry offers substantial advantages, there are several important considerations regarding its use in animal studies in high-containment settings. First, surgical techniques used for device implantation must be refined to improve device sensitivity and durability prior to entry into the high-containment facility, thereby minimizing the risk of mechanical or technical failures, as opportunities for repair or replacement may be limited or unavailable. Second, the technical infrastructure required to collect, transmit, and store telemetry data from high-containment laboratories to office environments for downstream analysis can present logistical and operational challenges. Third, the use of anesthesia can significantly alter telemetry-derived physiological parameters, particularly EEG, ECG, heart rate, and blood pressure for up to several hours, therefore, these data must be excluded from subsequent analyses (48–54). Consequently, collection of additional samples that require anesthesia, such as blood or cerebrospinal fluid, may not be feasible without compromising telemetry data. Collectively, these considerations must be addressed prior to study initiation to ensure data quality and operational feasibility.

The cynomolgus macaque model has been previously investigated to study various aspects of encephalitic alphavirus infection (27–42, 55–60). Infection via the aerosol route is the most reliable method for producing severe disease. Following aerosol exposure, infection with WEEV and EEEV results in lethal disease, whereas VEEV infection does not produce lethal disease. However, similar to WEEV and EEEV, VEEV can cause febrile illness, viremia, alterations in hematology and chemistry. The VEEV-IAB TrD strain has been the most frequently used in NHP studies investigating VEEV pathogenesis (28–30, 32, 34, 36–40). Virus stocks generated through serial passaging of the original isolate or derived from cDNA clones can cause disease in NHPs via aerosol route at doses ranging of ∼1.0 to 8.0 log_10_ PFU. Studies have utilized telemetry; however, its use has been limited and has primarily focused on temperature monitoring.

In the present study, we sought to expand the use of telemetry to investigate critical physiological parameters that would be routinely assessed following human infection in a clinical setting. Cynomolgus macaques were infected via the aerosol route with a cDNA clone derived VEEV-IAB TrD at 6.0 log_10_ PFU, and consistent with previous studies VEEV-IAB was able to cause disease. However, unlike studies we investigated respiration, activity, heart rate, blood pressure, disease signs, ECG, and EEG. Following VEEV-IAB infection, all parameters were altered, however the duration and magnitude differed substantially. The most consistent alterations were body temperature, behavior, and brain waves, all of which were disrupted within 1 to 2 dpi in all NHPs. Fever persisted through 7–9 dpi and resolved between 8–10 dpi. Behavioral alterations included reduced fluid consumption, disruption of circadian rhythm, as well as increased nighttime activity with a simultaneous decrease in sleep throughout 9 dpi. Brain waves, particularly beta and gamma waves, were altered through 28 dpi and did not return to baseline. In contrast, respiration, heart rate, blood pressure, and ECG parameters displayed an intermittent pattern during the first 10 dpi. These parameters exhibited daytime values near baseline, whereas nighttime values were considerably altered.

ECG alterations were assessed by measuring RR, PR, QRS duration and QTc interval. As expected, these intervals are dependent on heart rate, shortening with increased rates and lengthening with decreased rates. During 3 to 10 dpi, heart rate remained elevated in all NHPs resulting in the shortening of RR, PR, QRS duration and QTc interval duration. Surprisingly, during this period, NHPs exhibited cardiac abnormalities, with prolonged RR (NHP #2), PR (all NHPs), QRS duration (NHP #3), and QRS interval (all NHPs). These abnormalities suggest disruptions in cardiac electrical conduction and are consistent with ventricular arrhythmias. VEEV-IAB infection of neurons, reduced fluid intake, sleep disruption may contribute to ECG abnormalities.

VEEV-IAB infection induced distinct and sustained alterations in brain wave activity throughout the study. During week one, alterations in the slower waves (e.g., delta, theta, sigma, and alpha) were observed, ranging from -250% to +500%. In contrast during week two to four, considerable and sustained alterations were observed in beta and gamma waves, with alteration ranging from -250% to +4,800%. These changes were consistently more pronounced at nighttime. Notably, the pronounced alteration of slow wave activity associated with sleep during normal circadian rhythm cycle was observed in awake NHPs. While increased beta and gamma wave activity, normally associated with wakefulness in NHPs during normal circadian rhythm cycle, was observed in NHPs during nighttime, particularly during weeks three and four. Lastly, beta and gamma waves failed to return to baseline throughout the four-week study. Taken together, these findings highlight the profound and prolonged impact of VEEV-IAB infection on brain waves following aerosol exposure.

To investigate the mechanisms underlying the sustained alterations in brain waves, a comprehensive neuropathology study was performed on NHP brain tissues at 28 dpi. Minimal or no microscopic lesions, viral RNA or protein, were observed in all NHPs. The minimal lesions observed comprised primarily of gliosis or perivascular cuffing. Rare instances of viral RNA were detected in gliosis lesions in three of the four NHPs, however, these were difficult to detect. Similar minimal neuropathologic changes and limited viral RNA detection have been reported in a previous study of VEEV-IAB (32). Thus, the lack of virus/host induced brain pathology as well as the absence of viral RNA or proteins cannot explain the persistent alteration of brain wave activity.

One potential mechanism that may contribute to sustained alteration of the brain wave activity is traumatic brain injury (TBI). TBI is defined as a non-degenerative and non-congenital insult to the brain that results in temporary or permanent impairment of cognitive and physical functions (61). All four NHPs exhibited many signs associated with TBI such as disturbances in circadian rhythm, food/fluid consumption, inability to initiate or maintain a normal sleep pattern, decreased overall activity, and increased slow wave (delta, theta, and alpha) activity during wakefulness (61–64). This data strongly suggests that VEEV-IAB infection via the aerosol route can rapidly induce many features resembling severe TBI. In recent studies examining brain tissues of VEEV-IC infected cynomolgus macaques and serum samples from confirmed human VEEV-ID infection, found elevated levels of traumatic brain injury markers such as leukemia inhibitory factor (LIF), matrix metalloproteinase 9 (MMP-9), and glial fibrillary acidic protein (GFAP) (33, 65). These data support the hypothesis that VEEV infection can increase TBI markers and further studies are underway to investigate TBI markers in the brain.

The hallmark of acute infection in cynomolgus macaques following aerosol exposure to either EEEV or VEEV IAB infection at comparable doses is the development of fever. A recent study of EEEV infection reported temperature kinetics and magnitude are comparable to those observed in VEEV-IAB infection (42). However, the magnitude and duration of alterations in physiological parameters such as respiration, heart rate, blood pressure, ECG, and EEG differed substantially. One potential explanation of this difference may be due to the extent of virus replication and dissemination within brain regions comprising the autonomic nervous system.

ECG and EEG in cynomolgus macaques following VEEV-IC infection has been explored previously (31, 33). The ECG findings in this study, including RR, QT, and QRS intervals, were comparable to those reported for VEEV-IC. In contrast, the EEG results from this study are not comparable to the previous study. One potential explanation for the discrepancy may be due to the EEG lead placement to detect brain waves. The methodology for the lead placement in this study was extensively optimized in collaboration with Data Sciences International and Charles River to increase the sensitivity of the EEG signal.

Regulatory approval of medical countermeasures for Risk Group 3 and 4 agents will rely on the U.S. FDA Animal Rule (21 CFR 601.90), which allows pivotal efficacy studies in animal models to support licensure when human trials are not feasible. In this context, telemetry-based measurement of clinically relevant parameters offers several advantages for animal model development. First, it enables continuous, high-resolution monitoring of physiological endpoints that are directly translatable to human disease. Second, it supports the identification and quantification of multiple disease parameters, facilitating rapid refinement of animal models, and evaluation of countermeasures during natural outbreaks or bioterrorism events. This capability is particularly valuable for partially lethal or non-lethal pathogens such as western equine encephalitis virus (WEEV) and SARS-related coronaviruses. Third, the high sampling frequency of telemetry systems enables real-time assessment of treatment effects on a daily to hourly basis. Finally, telemetry may help identify potential adverse physiological effects of candidate countermeasures prior to human clinical evaluation. These advantages highlight the value of advanced telemetry in nonhuman primate models for studying high-consequence pathogens and support further investigation of its application in Risk Group 3 and 4 disease research.

In summary, we utilized state-of-the-art telemetry to investigate critical physiological parameters in the cynomolgus macaque model following VEEV-IAB aerosol exposure. All parameters measured including body temperature, behavior, respiration, heart rate, blood pressure, ECG, and EEG were altered within 1 to 2 dpi. The most altered and sustained parameters were body temperature, behavior, and EEG activity. Brains waves exhibited a marked and sustained disruption following infection and did not return to baseline in the 28-day study period. This is the first detailed disease course of VEEV-IAB in an NHP model, and the parameters identified will improve future animal model development and countermeasure evaluation.

## MATERIALS AND METHODS

### Virus and Cells

Venezuelan equine encephalitis virus subtype IAB Trinidad donkey strain was derived from a cDNA clone, rescued, and amplified in Vero-E6 cells (ATCC). The virus stock was deep sequenced to verify genomic sequence and to ensure purity. In addition, the stock was tested for endotoxin and mycoplasma.

### Ethics statement

Research was conducted under an Institutional Animal Care and Use Committee (IACUC) approved protocol in compliance with the Animal Welfare Act, Public Health Service Policy on Humane Care and Use of Laboratory Animals, and other federal statutes and regulations relating to animals and experiments involving animals. The facility where this research was conducted is accredited by the AAALAC International and adheres to the principles stated in The Guide for the Care and Use of Laboratory Animals, National Research Council, 2011.

### Nonhuman primate and study design

Four (1 males, 3 females) cynomolgus macaques (*Macaca fascicularis*) of Chinese origin ages 5 to 11 years and weighing 3 to 9 kg were obtained from Covance. Animals were implanted with an M11 and two M01 implants (Data Sciences International). Each device was dedicated to each hemisphere of the brain and was implanted at left and right scapula. The leads for each device were routed subcutaneously and surgically implanted at frontal and occipital lobes. The M11 implant was utilized to detect ECG, blood pressure, temperature, activity, and respiration. The two M01 implants were dedicated to detecting EEG activity in the brain with the electrodes of each implant placed at the frontal and occipital lobes. All NHPs were prescreened and determined to be negative for Herpes B virus, simian T-lymphotropic virus 1, simian immunodeficiency virus, simian retrovirus D 1/2/3, tuberculosis, *Salmonella* spp., *Campylobacter* spp., hypermucoviscous *Klebsiella* spp., and *Shigella* spp. NHPs were also screened for the presence of neutralizing antibodies to EEEV, VEEV IAB, and WEEV by plaque reduction neutralization test (PRNT_80_).

### Telemetry devices and data collection

The telemetry implantation in NHPs and data collection was completed as described previously (42).

### Establishing baseline for each physiological parameter

To establish a baseline of each parameter for individual animals, telemetry data was collected continuously for five days prior to infection. Data for each animal and parameter was utilized to generate a 0.5-hr interval by averaging up to 1,800 data points. Subsequently, a 48-point reference baseline for a 24-hr period was generated by averaging five previously calculated time-matched baseline values. Baseline 0.5-hr averages and standard deviations (SD) were generated. Daytime and nighttime were defined as 6 am to 6 pm and 6 pm to 6 am, respectively. All comparisons between pre- and post-infection were time matched. Following determination of baseline values, NHPs were infected with a target dose of 6.0 log_10_ PFU of VEEV-IAB via the aerosol route. The NHPs were observed for signs of disease and data for each parameter (temperature, activity, respiration and heart rates, blood pressure, ECG, and EEG) were obtained. Time-matched comparisons were made between pre-infection baseline and post-infection values.

### Aerosol infection

NHPs were exposed to the target inhalation dose of 6.0 log_10_ PFU of VEEV in the head-only Automated Bioaerosol Exposure System (ABES-II).The virus stock at 8.2 log_10_ PFU/mL was diluted to 8.0 log_10_ PFU/mL and utilized in the nebulizer. The inhalation infection was generated using a Collison Nebulizer to produce a highly respirable aerosol (flow rate 7.5±0.1 L/minute). The system generated a target inhalation of 1 to 3 μm mass median aerodynamic diameters determined by TSI Aerodynamic Particle Sizer. Samples of the pre-spray suspension and inhalation collected from the exposure chamber using an all-glass impinger (AGI) during the infection were analyzed by plaque assay to determine the inhaled PFU. The inhalation infection dose for each NHP was calculated from the minute volume determined with a whole-body plethysmograph box using Buxco XA software. The total volume of inhaled dose was determined by the exposure time required to deliver the estimated inhaled dose. Individual NHPs were infected successively in a head-only chamber under ABES-II.

### Post-exposure monitoring and score criteria

NHP observations began five days prior to aerosol exposure to obtain baseline data. Following aerosol infection, all NHPs were monitored daily via continuous 24-hr remote monitoring to limit room entries. Signs of disease were observed and a score for each NHP was determined by criteria described previously.

### Plaque assay

The virus stock and dose for each NHP were titrated on Vero-76 cells as described previously (42).

### Plaque Reduction Neutralization Test (PRNT_80_)

Serum samples from pre-bleed and 28 dpi were heat-inactivated at 56°C for 30 min. The PRNT_80_ assay was performed as described previously (42).

### Tissues processing and histopathology

Brain and spinal cord tissues were obtained from all four NHPs. Tissues from various brain and spinal cord regions including frontal cortex, amygdala, hippocampus, hypothalamus, thalamus, corpus striatum, mesencephalon, medulla oblongata, cerebellum, and spina cord regions (cervical, thoracic, and lumbar) were sectioned as described previously (57).

### *In situ* hybridization

*In situ* hybridization (ISH) was performed as described previously . ISH probe targeting the genomic RNA fragment of VEEV (7823– 8748 of KP282671.1) was designed and synthesized by Advanced Cell Diagnostics (Cat# 483101).

### Immunohistochemistry

Immunohistochemistry (IHC) was performed as described previously (57). A rabbit polyclonal anti-alphavirus antibody (Internal USAMRIID stocks) was used at a dilution of 1:6,000.

### Statistics

All comparisons between pre- and post-infection were time-matched. GraphPad Prism version 7.00 for Windows (GraphPad Software, La Jolla, California, USA) software was utilized for statistical analysis. Significant differences in each parameter for each animal were determined using paired t-test or one-way ANOVA, followed by Tukey’s test.

